# Predicting the future direction of cell movement with convolutional neural networks

**DOI:** 10.1101/388033

**Authors:** Shori Nishimoto, Yuta Tokuoka, Takahiro G Yamada, Noriko F Hiroi, Akira Funahashi

## Abstract

Image-based deep learning systems, such as convolutional neural networks (CNNs), have recently been applied to cell classification, producing impressive results; however, application of CNNs has been confined to classification of the current cell state from the image. Here, we focused on cell movement where current and/or past cell shape can influence the future cell fate. We demonstrate that CNNs prospectively predicted the future direction of cell movement with high accuracy from a single image patch of a cell at a certain time. Furthermore, by visualizing the image features that were learned by the CNNs, we could identify morphological features, e.g., the protrusions and trailing edge that have been experimentally reported to determine the direction of cell movement. Our results indicate that CNNs have the potential to predict the future cell fate from current cell shape, and can be used to automatically identify those morphological features that influence future cell fate.

## Introduction

Recent advances in microscope automation have enabled the acquisition of large numbers of bioimages. Several approaches to analyzing these images are proposed. One successful approach is machine learning, which has been used primarily in cell classification[1]. Cell classification using conventional machine learning proceeds in two steps. Firstly, hand-crafted image features are extracted for each cell from the image (e.g., using Scale-Invariant Feature Transform[2], Histograms of Oriented Gradients[3], or CellProfiler[4]). Secondly, these features are used to train a classification model (e.g., Support Vector Machine[5], Adaptive Boosting[6]). As a result, the performance of these classifiers relies heavily on the appropriateness of the hand-crafted features chosen empirically. Moreover, because feature extraction and classifier training are independent of each other, they cannot work together to identify and use discriminative information maximally.

In recent years, deep learning methods have been used to overcome this limitation of conventional machine learning methods. Deep learning methods, especially convolutional neural networks (CNNs), automatically learn feature representations from the raw pixels of cell images. Therefore, CNNs can avoid using hand-crafted features. Furthermore, CNNs are jointly optimized with these feature representations to predict the class for each cell image. For general visual recognition tasks, CNNs have substantially outperformed conventional machine learning methods with hand-crafted features[7, 8], and they have been applied successfully to biological imaging[9, 10]. In cell classification, use of CNNs has produced impressive results[11, 12, 13, 14, 15, 16, 17] : e.g., the classification of abnormal morphology in MFC-7 breast cancer cells[13], the classification of cervical cells in cytology images[15]; the identification of malaria-infected cells[16]; and the automatic classification of Hep-2 (human epithelial-2) cell staining patterns[17].

The above applications of CNNs have focused on classification of the current cell state from the image. However, recent studies have demonstrated that the current and/or past cell shape influences the future cell fate even after the original shape is lost. Akamura et al. (2016) demonstrated that the shape of a V2 neural progenitor cell affects the stochastic fate decision-making of the daughter cells even after the V2 cell loses the original geometry through mitotic rounding and division[18]. Kozawa et al. (2016) used Bayesian inference to predict cell division-timing based on progressive cell-shape changes[19]. Therefore, an interesting question is whether CNNs can be used to predict future cell fate based on current and/or past cell shape.

Here, we focused on dynamic cell movement as a model system of cell shape influencing future cell fate. In general, cell movement can be conceptualized as a cyclic process[20]. The cell movement cycle begins with the formation of protrusions by actin polymerization. Protrusions are stabilized by adhering to the extracellular matrix (ECM). These adhesions serve as traction sites for the movement as the cell moves forward over them. At the cell rear, the adhesions with ECM are disassembled. Then the cell rear, the trailing edge, contracts mainly due to the pulling force generated by myosin II. As a result, the cell moves towards the protrusions. The motility and shape of individual migrating cells are closely related[21]. Jiang et al. (2005) demonstrated that the polarity of cell shape, i.e., wide front and a narrow rear, biases the direction of cell movement (i.e., moving direction)[22]. Ghosh et al. (2004) demonstrated that protrusions formed locally by actin polymerization define the moving direction[23]. We therefore hypothesized that CNNs can learn which morphological features influence moving direction and thus predict the future moving direction from cell shape at a certain time.

In general, it is not clear how and why CNNs arrive at a particular prediction decision, although several groups have attempted to interpret CNN predictions and propose possible methods for explaining CNNs decisions[24, 25, 26, 27]. These methods can visualize the features of the image that contribute to the CNN predictions. Application of these methods is expected to identify morphological features that influence moving direction.

Here, we demonstrate that CNNs prospectively predict the future direction of cell movement with high accuracy from a single image patch of a cell at a certain time. Furthermore, to reveal how and why CNN models can predict future moving direction, we visualized the features of the cell images that were learned by the CNN models and contributed to their prediction: e.g., the protrusions and trailing edge.

## Results

### Training and validation of CNNs for predicting the future direction of cell movement

To illustrate that CNNs can predict cell fate based on current cell shape, we set out to construct a CNN model that predicts the future direction of cell movement from a single image patch of a cell at a certain time. Firstly, we prepared image datasets from time-lapse phase-contrast microscopic images of migrating NIH/3T3 cells and U373 cells, respectively (see Materials and Methods). We manually tracked the positions of the migrating cells. Then, each cell in each frame was annotated with the moving direction: i.e., toward upper right, upper left, lower left, or lower right. The annotation was determined based on the displacement at the time the net displacement exceeded the average diameter of NIH/3T3 cells (18 *µ*m^*^). For each annotated cell in each frame, we created an image patch of 128*×*128 pixels centered on its position coordinate. The NIH/3T3 dataset comprised 785 image patches for 40 cells (Table 1); the U373 dataset comprised 795 image patches for 12 cells (Table 1).

**Table 1:** Number of image patches per moving direction

| Moving direction | Number of image patches<br>in the NIH/3T3 dataset | Number of image patches<br>in the U373 dataset |
| --- | --- | --- |
| upper right | 179 | 165 |
| upper left | 188 | 253 |
| lower left | 242 | 96 |
| lower right | 176 | 281 |

We used these datasets to train and test CNN models for predicting the direction of cell movement. In CNN models, input image patches are processed through multiple convolutional layers and max-pooling layers. In convolutional layers, trainable sets of filters are applied at different spatial locations using a stride of a certain size, thereby extracting features associated with moving direction as feature maps. In max-pooling layers, feature maps are down-sampled by computing the maximum value of a feature map over a region, which reduces variance and increases translational invariance[28]. After repeating the processing through convolutional layers and max-pooling layers, two fully connected layers are used for prediction. Our CNN models arrange 14 layers into eight convolutional layers, four max-pooling layers, and two fully connected layers, consisting of 162,704 trainable parameters in total (more detail in Material and Methods, network architecture shown in Fig. 1 and Table 2).

**Figure 1:**
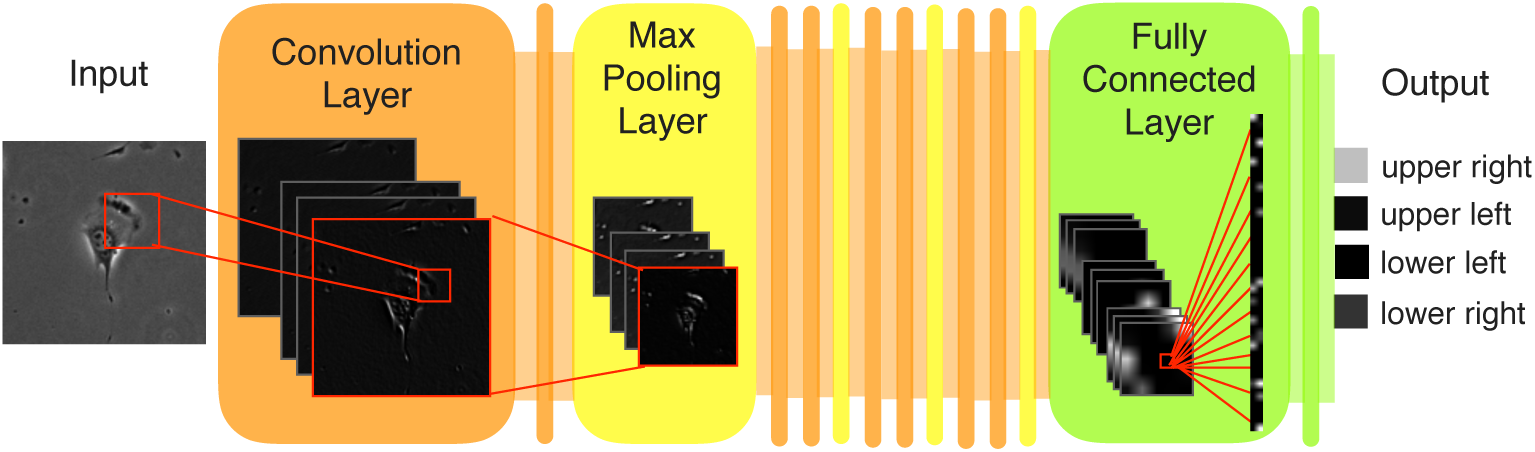
Architecture of CNN models that predict the future direction of cell movement using a single image patch of a cell at a certain time. In the flowchart, an image patch of a NIH/3T3 cell annotated on the upper right (Input) is presented to a CNN model. The input is processed through a series of repeating convolutional layers (orange) and max-pooling layers (yellow). In the convolutional layer, the activation images illustrate extracted feature maps of the sample image patch (Input). The red boxes and lines illustrate the connections within the CNN model. After repeating processing through convolutional layers and max-pooling layers, fully connected layers are used for prediction (green). The network output (Output) represents the distribution over four moving directions.

**Table 2:** CNN model-relevant hyperparameters

| Layer | Type | Description |
| --- | --- | --- |
| 1 | Convolution | Filter size = $3 \times 3$, Number of filters = 16, Stride size = 1 |
| 2 | Convolution | Filter size = $3 \times 3$, Number of filters = 16, Stride size = 1 |
| 3 | Max-pooling | Filter size = $2 \times 2$, Stride size = 2 |
| 4 | Convolution | Filter size = $3 \times 3$, Number of filters = 16, Stride size = 1 |
| 5 | Convolution | Filter size = $3 \times 3$, Number of filters = 16, Stride size = 1 |
| 6 | Max-pooling | Filter size = $2 \times 2$, Stride size = 2 |
| 7 | Convolution | Filter size = $3 \times 3$, Number of filters = 32, Stride size = 1 |
| 8 | Convolution | Filter size = $3 \times 3$, Number of filters = 32, Stride size = 1 |
| 9 | Max-pooling | Filter size = $2 \times 2$, Stride size = 2 |
| 10 | Convolution | Filter size = $3 \times 3$, Number of filters = 64, Stride size = 1 |
| 11 | Convolution | Filter size = $3 \times 3$, Number of filters = 64, Stride size = 1 |
| 12 | Max-pooling | Filter size = $2 \times 2$, Stride size = 2 |
| 13 | Fully connected | Number of neurons = 100 |
| 14 | Fully connected | Number of neurons = 4 |

To assess the generalization performance of CNN models, we trained and tested them in 4-fold cross-validation using the NIH/3T3 and U373 datasets. We evaluated the performance of trained CNN models by calculating the average classification accuracy (ACA) and the mean class accuracy (MCA) (see Materials and Methods); the results are presented below as means *±*standard deviation (n = 4 folds). While the expected value of ACA and MCA, if the data were randomly predicted, are 25%, our CNN models achieved high ACA of 87.3% *±* 3.0% and high MCA of 85.9% *±* 3.1% for the NIH/3T3 dataset. Our CNN models also achieved high ACA of 89.1% *±* 0.4% and high MCA of 86.7%*±* 1.1% for the U373 dataset.

### Visualizing image features learned by the CNN models

Next, to identify the morphological features influential in the prediction of CNN models, we quantified and visualized the image features learned by the CNN models. Firstly, we used guided backpropagation (GBP)[27], which visualizes the local feature of the input image that most strongly activates a CNN’s particular neuron (see Material and Methods). For each dataset, we presented each test image to the trained CNN models. Then, GBP was applied to a single maximum activation in each feature map of the last convolutional layer, producing local features of the input image patch. As a result, local features corresponding to the protrusion and the trailing edge contracting in the moving direction were identified for both the NIH/3T3 and U373 datasets (lower parts of each image group, Fig. 2). Next, we quantified and visualized global features of the image patches contributing to the CNN prediction of the moving direction by using deep Taylor decomposition (DTD)[26]. DTD quantifies the degree of contribution to the CNN prediction (relevance) of the input image patch (see Materials and Methods) on a pixel-by-pixel basis. For each dataset, we presented each test image to the trained CNN models. Then, by applying DTD to the prediction result, pixel-wise relevances were calculated. We visualized these pixel-wise relevances as heatmaps (Fig. 2, lower right). For both the NIH/3T3 and U373 datasets, pixels corresponding to the protrusion and trailing edge simultaneously contributed to the prediction of the CNN models.

**Figure 2:**
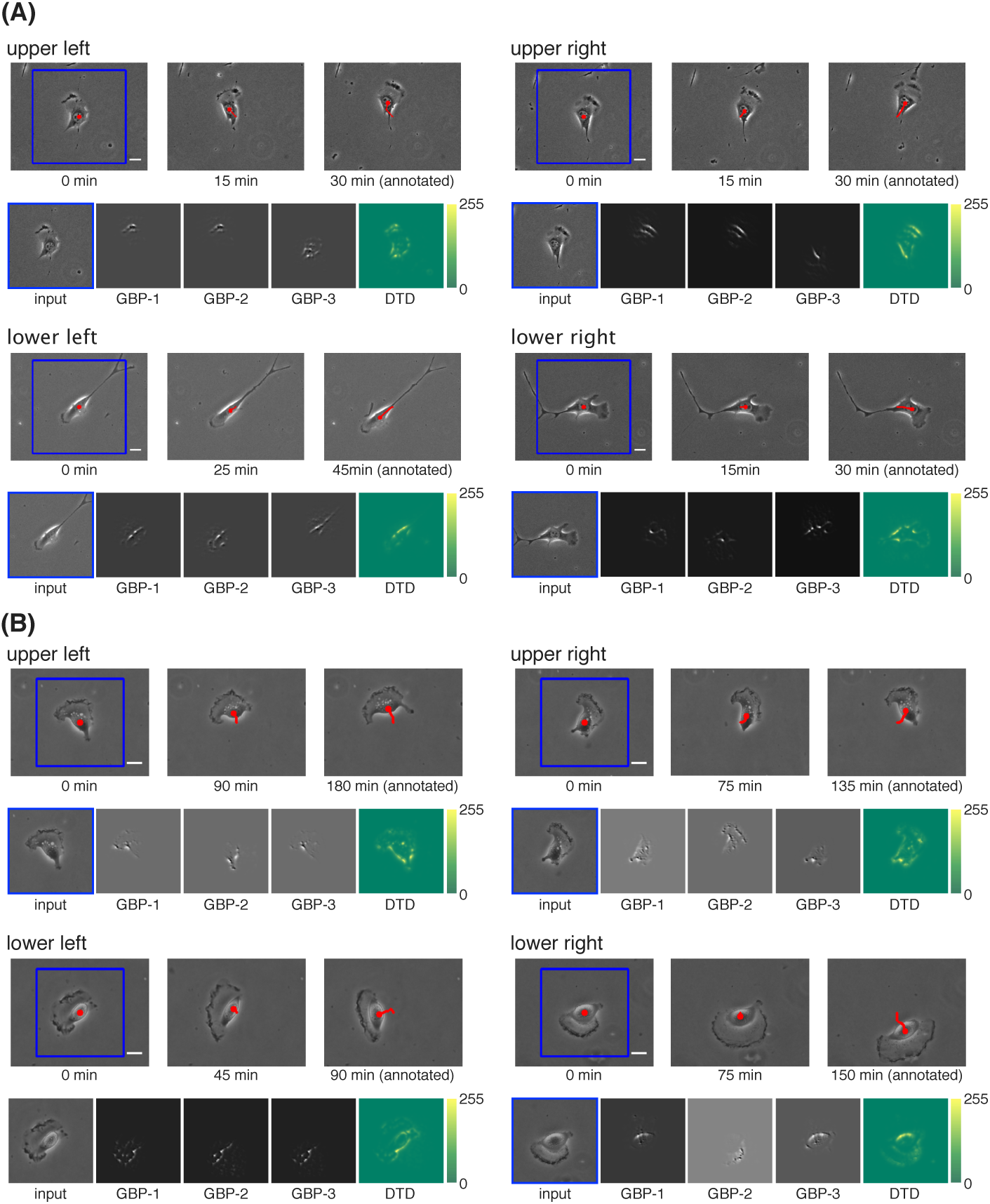
Visualized image features learned by the CNN models. (A) NIH/3T3 dataset. (B) U373 dataset. For each moving direction, each group of images shows exemplary results (i.e., those for a correctly predicted test image patch). The upper row of each group of images comprises—from left to right—the frame corresponding to the input image patch, the frame imaged in the middle between the left frame and the right frame, and the frame when the moving direction was annotated (scale bars, 20 *µ*m). The time under each frame shows the elapsed time since the leftmost frame was imaged. The blue bounding box indicates the area corresponding to the input image patch. The red dot indicates the position of the cell obtained by manual tracking. The red line indicates the trajectory of cell movement starting from the position of the cell at 0 min. The lower row of each group of images comprises—from left to right—the input image patch, the local features visualized by GBP for the feature maps whose maximum activations were top three, and the heatmap of pixel-wise relevance calculated by DTD.

## Discussion

Previous applications of CNNs to cell classification have focused on classification of the current cell state from an image[11, 12, 13, 14, 15, 16, 17]. In contrast, here, we focused on images of dynamic cell movement and demonstrated that CNNs can prospectively predict the future direction of cell movement with high accuracy.

Average speed and directionality are important parameters that define cell motility[29]. Both these parameters were significantly different between the NIH/3T3 and the U373 datasets (Supplementary Fig. 1). The apparent morphology of migrating cells also differed qualitatively between the two datasets: e.g., the protrusions were prominent and broad in U373 cells compared with NIH/3T3 cells (Fig. 2, Movie 1 and 2 in Supplementary Material). Despite these differences, our CNN models achieved the same degree of prediction accuracy for both cell types. This might be because our CNN models could learn the morphological features, such as the protrusion and the trailing edge (Fig. 2), which are known to be characteristic features of cell movement regardless of cell type or substrate, except for some specific cell types[20, 30].

Because cell classification using conventional machine learning is built upon the extraction of hand-crafted features[1, 2, 3, 4], the features affecting the classification result depend on the choice of hand-crafted features. These handcrafted features are abstract representations of the cell image and are therefore diffcult to intuitively relate to visual inspection of the cell image. In contrast, because CNNs automatically and jointly learn optimal feature representations and the prediction task directly from the raw pixels of the cell image, the features affecting the prediction result do not depend on the feature selection and are optimized for the prediction. Furthermore, because these features can be visualized on the cell image by using the approach used in this study, they can be intuitively related to visual inspection of the cell image. Here, we identified morphological features, such as protrusions and trailing edge, which are known experimentally to influence the moving direction[for review, see [20]]: for instance, protrusions generated locally by actin polymerization determine the moving direction[23], and the position of the trailing edge influences the moving direction via stress fiber organization[31]. Our results indicate that such important morphological features that can influence cell fate can be identified automatically, by using CNNs to predict cell fate and then visualizing the image feature(s) contributing to this prediction.

In conclusion, we focused on dynamic cell movement as a model system in which cell shape influences cell fate. Our findings indicate that use of CNNs can enable not only the prediction of the cell fate but also the identification of morphological features that can influence the cell fate. Based on these findings, we believe that our approach could be applied to the analysis of other cell mechanisms where cell shape can influence future processes, such as cell division[19] and differentiation[32]. This approach could reveal unknown biological phenomena and be useful for pathologic and/or clinical diagnosis.

## ACKNOWLEDGEMENTS

We thank the Doi laboratory at Keio University for providing NIH/3T3 fibroblasts. We thank Takumi Hiraiwa from the Department of Biosciences and Informatics at Keio University for assistance with experiments. We are also grateful to Carlos Ortiz-de-Solorzano from the Center for Applied Medical Research (Pamplona, Spain) for providing the dataset used in the ISBI cell tracking challenge 2015. The research was funded by a JSPS KAKENHI Grant (Number 16H04731).

## Author contributions

A.F. initiated the project; A.F. and N.H. supervised the project; S.N. prepared time-lapse phase-contrast microscopy images of NIH/3T3 cells, created the datasets, developed the convolutional neural network approach, analyzed the data, and wrote the manuscript with Y.T. and T.Y.; A.F. and Y.T. gave technical advice on analysis; N.H. advised from a biological point of view. All authors commented on the manuscript.

## Materials and Methods

### Time-lapse phase-contrast microscopic images

To create datasets for training CNN models, we prepared time-lapse phasecontrast microscopic images of cell movement in different ways for each dataset; details are provided below.

### NIH/3T3 dataset

NIH/3T3 fibroblasts were plated on glass-based dishes precoated with 5 *µ*g/cm^2^ fibronectin at a density of 500 cells/cm^2^. After overnight incubation, cell movements were monitored with an inverted microscope (IX81, Olympus) equipped with an on-stage incubation chamber that maintained the temperature at 37^*°*^C and the CO_2_ concentration at 5%, using a 20*×* objective (0.45 numerical aperture). Phase-contrast images were collected with CCD video cameras (ORCAFlash4.0 or ORCA-R2; Hamamatsu) at 5-min intervals, digitized and stored as image stacks by using Micro-Manager software (Open Imaging).

### U373 dataset

Time-lapse phase-contrast microscopic images of glioblastoma–astrocytoma U373 cells were obtained from the dataset used in the ISBI (International Symposium on Biomedical Imaging) cell tracking challenge 2015[33, 34]. U373 cells moving on a polyacrylamide substrate were monitored with a microscope (Nikon) using a 20*×* objective (0.5 numerical aperture), and phase-contrast images were acquired at 15-min intervals.

### Annotation of the moving direction

For each dataset, we manually tracked the positions of migrating cells by using the Manual Tracking plugin of ImageJ (National Institutes of Health). For each cell in each frame, we annotated one of the four moving directions according to the value of displacement (Δ*x*, Δ*y*) at the time the net displacement 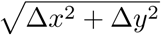 (the distance between the initial and final positions) exceeded the average diameter of NIH/3T3 cells (18 *µ*m^†^)(Fig. 3). The four moving directions were defined as follows: (i) Δ*x≥* 0, Δ*y≤* 0 = upper right; (ii) Δ*x <* 0, Δ*y≤* 0 = upper left; (iii) Δ*x <* 0, Δ*y >* 0 = lower left; and (iv) Δ*x≥* 0, Δ*y >* 0 = lower right. Regarding the NIH/3T3 dataset, the net displacement was evaluated at 15-min intervals according to the shooting interval of the U373 dataset. For each annotated cell, we cropped the image to a bounding box of 256*×*256 pixels for the NIH/3T3 dataset or 170*×*170 pixels for the U373 dataset, centered on the cell *x, y* coordinates. Cropped image patches were resized to 128*×*128 pixels for each dataset.

**Figure 3:**
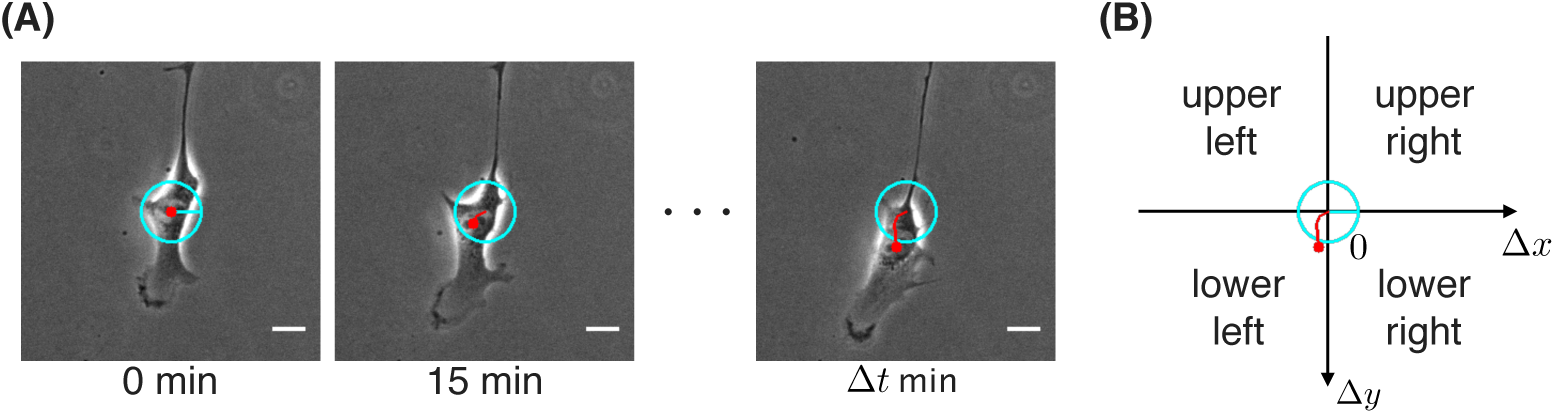
Annotation of the moving direction. (A) Exemplary time-lapse images of a migrating cell (scale bars, 20 *µ*m). The time under each frame shows the elapsed time since the leftmost frame (annotation target) was imaged. Δ*t* is the time when the net displacement 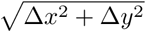 exceeded the average diameter of NIH/3T3 cells. The red dot indicates the position of the cell obtained by manual tracking. The red line indicates the trajectory of cell movement starting from the position of the cell at 0 min. The radius of the cyan circle is the average diameter of NIH/3T3 cells. (B) Annotating one of the four moving directions. According to the value of cell displacement (Δ*x*, Δ*y*) at the time Δ*t*, the moving direction was annotated as shown in the figure. The red line and the cyan circle are the same as those of the frame at Δ*t* min in (A).

### CNNs for predicting the future direction of cell movement

To prospectively predict the future direction of cell movement from a single image patch of a cell at a certain time, we used CNN models[35, 36] with the architecture shown in Fig. 1 and Table 2. Each of our CNN models comprised consecutive convolutional layers, max-pooling layers, and fully connected layers. The last fully connected layer consisted of four neurons, with each neuron corresponding to the future direction of cell movement. We applied the softmax function to the activations of the last fully connected layer to produce a distribution over four moving directions. Each convolutional layer and the first fully connected layer was followed by a nonlinear activation function. We used rectified linear units, which have been shown to introduce nonlinearities without suffering from the vanishing gradient problem[37]. To enable the CNN models to predict the future moving direction for image patches with various intensity values, the pixel values of each input image patch were normalized to within the range of 0 to 1 by first subtracting the minimum intensity value of the image patch, and then dividing by the maximum intensity value of the resulting image patch[17].

### Training and validation of CNN models

We trained CNN models and tested them in 4-fold cross-validation. At each fold, we used a softmax loss function and trained the CNN model for 50 epochs, using stochastic gradient descending with stratified batches of 32 images. To reduce the influence of imbalanced moving directions in the datasets, we multiplied the loss function by the class weight expressed by the following equation:

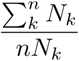

where *n* is the number of classes (moving directions), and *N*_*k*_ is the number of training data of class *k*. We initialized all weights in a CNN model by using the HeNormal algorithm, which sets the weights of each layer according to a scaled Gaussian distribution whose standard deviation corresponds to the input size of each layer[38]. We used standard values for the base learning rate (0.01) and momentum (0.9). For each epoch, we evaluated the prediction accuracy of a CNN model using test data. The evaluation criteria were the ACA and the MCA: ACA is the overall correct prediction rate of all test data, and MCA is the average of the per-class accuracies defined as follows:

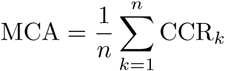

where CCR_*k*_ is the prediction accuracy of class (moving direction) *k* and *n* is the number of classes. If the MCA was not improved for the best score in the previous epoch, the learning rate was decreased by multiplying by 0.9. To prevent the CNN models from overfitting to training data, we (a) trained the models using vertical and horizontal reflections of the image patches as well as rotations by 90^*°*^, 180^*°*^, and 270^*°*^, in addition to the original training data; and (b) applied dropout[39] to the first fully connected layer (dropout rate, 0.5). For each fold, we used the best score of each criterion for 50 epochs as the prediction accuracy of the models.

For the following analysis of image features learned by CNN models, we used those models with the best MCA for the test data at each fold. We implemented and trained the CNN models by using Chainer[40], which is open-source software for machine learning.

### Visualizing image features learned by CNN models

We quantified and visualized features learned by the CNN models as described in the Results. GBP[27] was used to visualize local features of input image patches specific to the annotated moving direction. In this method, an input image is presented to a CNN, and high-level feature maps are computed throughout the layers. Typically, a single neuron activation is left non-zero in the high-level feature map. The single non-zero activation is back-propagated to input pixel space. Finally, the part of the input image that is most strongly activating the single non-zero neuron (i.e., the part that is most discriminative) is reconstructed. For each dataset, we presented each test image patch to trained CNN models, and a single maximum activation in each feature map of the last convolutional layer was back-propagated to input pixel-space, resulting in producing local features of the input image.

To quantify and visualize global features of input image patches contributing to the CNN prediction of moving direction, we used DTD[26], which can be summarized as follows. As a premise, each neuron of a CNN is viewed as a function that can be expanded and decomposed on its input variables. A CNN prediction of the moving direction is obtained by forward-propagation of input pixel values *{x_p_}* and is encoded by the output neuron *x*_*f*_. The output neuron is assigned a relevance score *R*_*f*_ = *x*_*f*_ representing the total evidence for the specific moving direction. Relevance is then decomposed and back-propagated from the top layer down to the input. As a result, the pixel-wise relevance (i.e., the degree of contribution to the CNN prediction) is calculated for each pixel of the input image. The pixel-wise relevances can be visualized as a heatmap. For each dataset, we presented each test image patch to trained CNN models, and DTD was applied to the CNN prediction output.

### Data availability

The image datasets used to train the CNN models, the raw time-lapse phase contrast images, metadata text files of Micro-Manager, and tracking results are available using bash scripts in: https://github.com/funalab/PredictMovingDirection.

### Code availaility

The code for performing the experiments is available at: https://github.com/funalab/PredictMovingDirection.

## Supplementary Material

**Supplementary Figure 1. Motility values calculated for the NIH/3T3 and U373 datasets.** (A) Boxplot of the average speed of the cell, *v*. (B) Boxplot of the directionality of the cell, *k* (*n* = 785 cell images in the NIH/3T3 dataset and 795 cell images in the U373 dataset). P-value is from two-sided Mann-Whitney rank test. For each image in the datasets, we first measured the time required until the net displacement Δ*r* exceeded the average diameter of NIH/3T3 cells, which is the time interval Δ*t* until the moving direction was annotated. Then, we calculated the total distance ∑Δ*d* traveled by the cell in the time interval Δ*t*. Regarding the NIH/3T3 dataset, The directionality was Δ*r* and movement distance Δ*d* were calculated at 15-min intervals according to the shooting interval of the U373 dataset. The average speed was calculated by the equation *v* =∑Δ*d/*Δ*t*[22]. The directionality was calculated by dividing the net displacement the net displacement Δ*r* by the total distance ∑Δ*d*[22]. used to measure how often the cell tended to turn. Cells that frequently make turns will yield a *k* value close to 0, whereas cells that persistently move along one direction will yield a *k* value close to 1.

**Movie 1. Time-lapse phase-contrast images of migrating NIH/3T3 cells used to create the NIH/3T3 dataset (AVI).** Images were acquired as described in Materials and Methods.

**Movie 2. Time-lapse phase-contrast images of migrating U373 cells used to create the U373 dataset (AVI).** Images were acquired as described in Materials and Methods.

## Footnotes

* http://bionumbers.hms.harvard.edu/bionumber.aspx?id=108905

† http://bionumbers.hms.harvard.edu/bionumber.aspx?id=108905

